# BIOTIN ATTACHMENT DOMAIN-CONTAINING proteins, inhibitors of ACCase, are regulated by WRINKLED1

**DOI:** 10.1101/634550

**Authors:** Hui Liu, Zhiyang Zhai, Kate Kuczynski, Jantana Keereetaweep, Jorg Schwender, John Shanklin

## Abstract

WRINKLED1 (WRI1) is a transcriptional activator that binds to AW boxes in the promoters of many genes from central metabolism and FA synthesis, resulting in their transcription. BIOTIN ATTACHMENT DOMAIN-CONTAINING (BADC) proteins are homologs of BIOTIN CARBOXYL CARRIER PROTEIN (BCCP) that lack a biotin-attachment domain and are therefore inactive. In the presence of excess FA, BADC1 and BADC3 are primarily responsible for the observed long-term irreversible inhibition of ACETYL-COA CARBOXYLASE (ACCase), and consequently FA synthesis. Purified WRI1 bound with high affinity (*Kd*s in the low nanomolar range) to canonical AW-boxes from the promoters of all three *BADC* genes. Consistent with this observation, the expression of *BADC1, BADC2 and BADC3* genes and BADC1 protein levels were reduced in *wri1-1* relative to wild type (WT), and *BADCs* gene expression and BADC1 protein levels also were elevated upon *WRI1* overexpression. The double mutant *badc1badc2* phenocopied *wri1-1* with respect to both reduction in root length, and elevation of indole-3-acetic acid-Asp (IAA-Asp) levels relative to WT. Overexpression of *BADC1* in *wri1-1* decreased its IAA-Asp and partially rescued its short-root phenotype demonstrating a role for BADCs in seedling establishment. That WRI1 positively regulates genes encoding both FA synthesis and BADCs i.e., conditional inhibitors of FA synthesis, represents a coordinated mechanism to achieve lipid homeostasis in which plants couple the transcription their FA synthetic capacity with their capacity to biochemically downregulate it.

**One sentence summary:** WRI1 regulates genes encoding both fatty acid synthesis and inhibitors of FA synthesis (BADCs), creating a lipid homeostatic mechanism in which the transcription of FA synthetic capacity is coordinated with the capacity to biochemically downregulate FA synthesis.

## Introduction

Lipids are primary metabolites in cells, acting as structural components of cell membranes, energy-dense storage compounds and cell signaling molecules. Fatty acids (FAs) are major components of lipids and triacylglycerols (TAGs) which are storage lipids that accumulate mostly in oil bodies in plant seeds (Li-Beisson et al., 2013). De novo synthesis of FAs occurs in the plastid via a well-established pathway (Ohlrogge and Browse, 1995; Rawsthorne, 2002), In Arabidopsis, transcription factors (TF): LEAFY COTYLEDON1 (LEC1) and LEC2, FUSCA3 (FUS3), and ABSCISIC ACID INSENSITIVE3 (ABI3), serve as the master regulators of embryogenesis and seed maturation (Parcy et al., 1997; Lotan et al., 1998; To et al., 2006) and also regulate accumulation of TAG (Kagaya et al., 2005; Mendoza et al., 2005; Braybrook et al., 2006). WRINKlED1 (WRI1), an APETALA2 (AP2)-type transcription factor, functions downstream of LEC1 and LEC2 (Baud et al., 2007; Mu et al., 2008) and is a master regulator of FAs synthesis because seeds of the *wri1* mutant show an 80% reduction in TAGs compared to wild type (WT) (Focks and Benning, 1998). More than 20 *WRI1* target genes have been identified by comparing gene expression in WT with that of *wri1* and *WRI1* overexpression-lines, and the use electrophoretic mobility shift assay (EMSA) confirmed WRI1-gene promoter binding. Based on promoter sequence comparisons [CnTnG](n)_7_[CG] was identified as the consensus AW-binding site. WRI1 target sequences are found upstream of genes coding for enzymes involved in glycolysis (sucrose synthase (SUS2), pyruvate kinase, and pyruvate dehydrogenase), Glu6P and PEP plastidial transporters (Glu6P/phosphate translocator and PEP/phosphate translocator), subunits of acetyl-CoA carboxylase (ACCase) (biotin carboxyl carrier protein 2 (BCCP2), biotin carboxylase (BC) and carboxyltransferase (CTα)), fatty acid synthesis (malonyl-CoA:ACP malonytransferase, ketoacyl-ACP synthase, hydroxyacyl-ACP dehydrase, enoyl-ACP reductase, acyl-carrier proteins, oleoyl-ACP thioesterase (FatA)) and genes involved in lipoic acid synthesis, a cofactor of pyruvate dehydrogenase (Ruuska et al., 2002; Baud et al., 2007; Maeo et al., 2009; Fukuda et al., 2013; Li et al., 2015). Interestingly, genes involved in TAG synthesis and oil body assembly such as diacylglycerol acyltransferase 1 (DGAT1) that catalyzes the final step of TAG synthesis and oleosins and caleosins appear not to be regulated by WRI1, though some species-specific variation has been reported (Maeo et al., 2009; Grimberg et al., 2015). By characterizing gene expression of subunits of ACCase, it was shown that the distance between the AW site and the translational initiation site (TIS) strongly influences the function of the AW box (Fukuda et al., 2013) in that the majority of AW sites fall within 200bp of the TIS in *bona fide* WRI1 target genes (Maeo et al., 2009; Fukuda et al., 2013).

A class of proteins annotated as biotin attachment domain-containing (BADC) (Olinares et al., 2010) have been recently identified to be inhibitors of FAs synthesis (Salie et al., 2016; Keereetaweep et al., 2018). BADC proteins are homologs of BCCP, sharing 25-30% sequence similarity, but they lack a biotin attachment site, rendering them inactive. All three Arabidopsis BADC isoforms interact with BCCP. Displacement of functional BCCP subunits in ACCase by inactive BADC subunits was believed to reduce the ACCase catalytic activity (Salie et al., 2016). ACCase is subject to feedback-regulation upon exposure to exogenously supplied fatty acids in the form of Tween esters (Tween-80 containing predominantly oleic acid; 18:1). Short-term exposure results in reversible inhibition of ACCase, whereas longer-term exposure results in irreversible inhibition (Andre et al., 2012). Oleoyl-ACP was shown to mediate the reversible phase of inhibition (Andre et al., 2012), whereas BADC1 and BADC3 are primarily responsible for the phase of irreversible inhibition (Keereetaweep et al., 2018).

In this work we present evidence that the expression of all three *BADC* genes are under the control of the WRI1 transcription factor. The short-root phenotype and elevated conjugated IAA levels common to the *wri1-1* and the *badc1badc2* double mutant led us to hypothesize that BADC-deficiency is responsible for the observed short-root phenotype. Restoration of near-WT root length upon overexpression of BADC1 in both *wri1* and *badc1badc2* is consistent with a yet-to-be identified role for BADCs in seedling development.

## Results

### *badc1 badc2* double mutant shows reduced primary root growth and fewer lateral roots than WT

Seeds of the *badc1 badc2* double mutant generated in our previous study, contained 17% and 23% increases in total fatty acid and TAG accumulation, respectively compared to WT (Keereetaweep et al., 2018). When grown on vertical MS plates, *badc1 badc2* exhibited a reduction in primary root length by 62% and a decrease in lateral root number by 39% compared with WT (Fig. 1A, 1B and 1C). In contrast, root growth of *badc1* or *badc2* single mutants showed no differences from that of WT. A complementation experiment was performed in which either *BADC1* or *BADC2* was expressed under the control of the constitutive 35S promoter in the *badc1 badc2* mutant. More than 10 independent lines were generated for each of *BADC1* and *BADC2* constructs into *badc1 badc2* (SFig. 1). All transgenic lines demonstrated restored root growth to varying degrees, indicating the involvement of *BADC1* and *BADC2* in seedling development (Fig. 1D and 1E). Because the *wri1* mutant (*wri1-1*) was recently shown to have short-root phenotype during seedling establishment (Kong et al., 2017), we compared the growth of *badc1 badc2* and *wri1-1* under the same conditions and show that their phenotypes are visually indistinguishable (SFig. 2).

**Figure 1.**
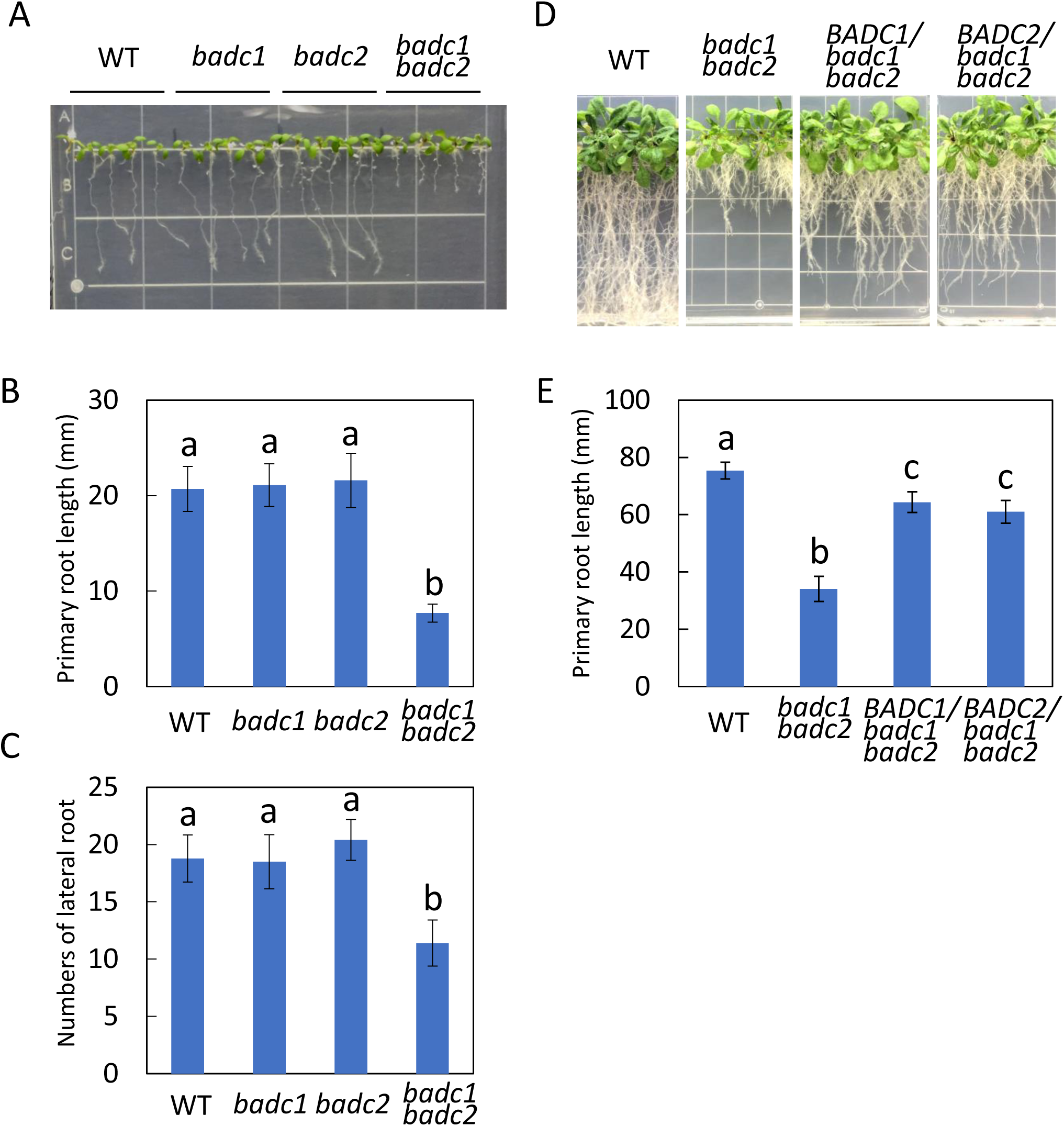
*badc1 badc2* double mutant shows reduced primary root growth and less lateral roots number than wild type (WT). (A) 9-day-old WT, *badc1, badc2*, and *badc1 badc2* seedlings grown vertically on 1/2MS media supplemented with 1% of sucrose. (B) Primary root length measurement of each genotype in (A). (C) Lateral root numbers for each genotype in (A). Values in (B) and (C) represent means±SD from 10 individual plants for each genotype. (D) Roots growth and (E) primary roots length of 20-day-old plants of WT, *badc1 badc2*, and representative transgenic *badc1 badc2* overexpressing *BADC1* or *BADC2* lines. Values in (E) are means±SD from 10 individual plants for each indicated genotype. In the figure, Levels indicated with different letters above histogram bars are significantly different (Student’s T test for all pairs of genotypes, P<0.01). Data in this figure is a representative of 3 independent repetitions.

**Figure 2.**
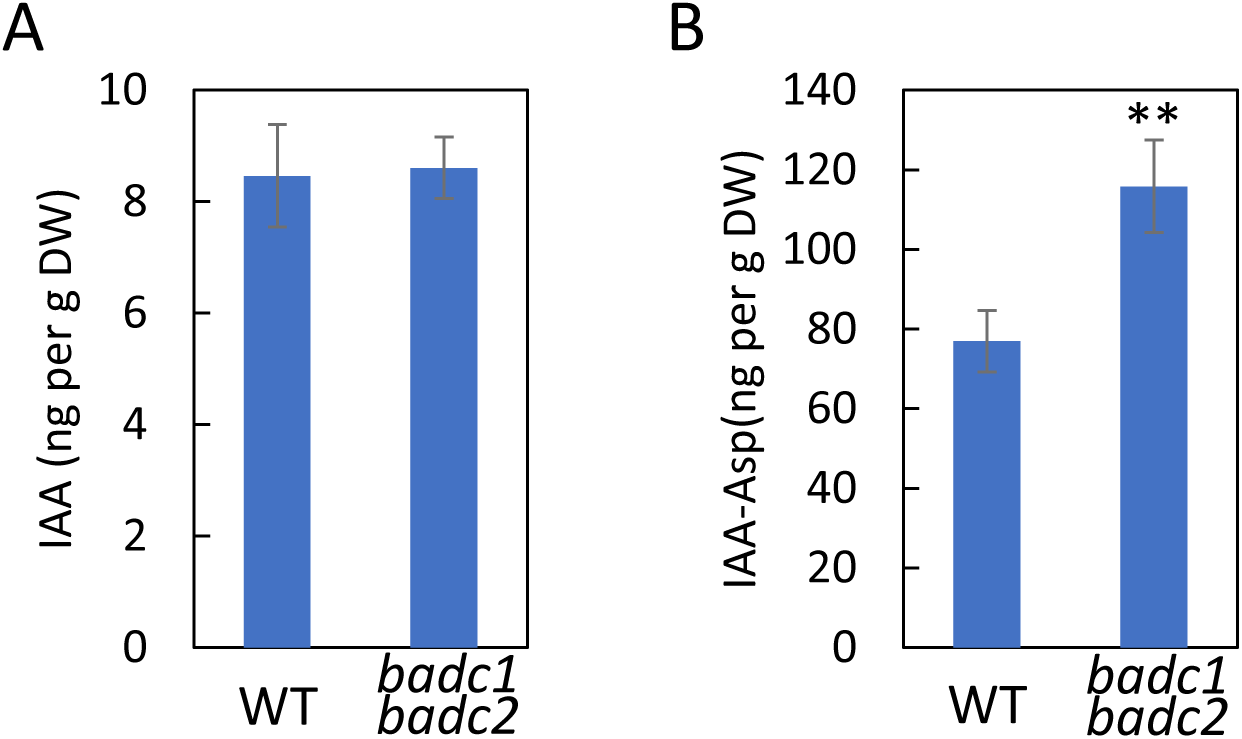
IAA-Asp content in *badc1badc2* double mutant is significantly higher than that in WT. (A) Quantification of IAA in 7-day-old seedling of WT and *badc1badc2*. (B) Quantification of IAA-Asp in WT and *badc1badc2*. Values in this figure are means ±SD (n=4) for each sample of 30 seedlings. Asterisks denote statistically significant difference from WT (Student’s T test, **, P<0.01). Data in this figure is a representative of 2 independent repetitions.

### Levels of conjugated auxin were elevated in *badc1 badc2* plants compared to those of wild-type

In *wri1-1* the levels of conjugated auxin, i.e., indole-3-acetic acid (IAA)-Asp, were reported to be significantly elevated compared to WT (Kong et al., 2017). The similarity between the root phenotypes of *badc1 badc2* and *wri1-1* during seedling establishment prompted us to measure the levels of several growth hormones. As shown in Fig. 2, IAA-Asp levels in 7-day-old seedlings showed a highly significant approximately 50% increase relative to levels found in WT. In contrast, the levels of other plant hormones i.e., ABA, JA and SA were not significantly different between *badc1 badc2* and WT (Table S1).

### WRI1 interacts with AW boxes from the promoters of *BADCs*

That *badc1 badc2* mimics *wri1-1* in both its short-root phenotype and IAA-Asp content led us to hypothesize that *BADCs* are under transcriptional control of WRI1. Sequence analysis identified a canonical AW-box consensus sequence within 200bp upstream of the translational initiation site (TIS) in the promoters of all three *BADC* isoforms (Fig. 3A and Table S2). To test whether WRI1 binds directly to these AW boxes, microscale thermophoresis (MST) assays were performed, in which a 28bp dsDNA fragment (Table S2), containing each of AW boxes from *BADCs* was titrated against purified WRI1 DNA binding domain (AA_58-240_) fused with GFP (SFig. 3). As shown in Fig. 3B and 3C, WRI1 exhibited low equilibrium dissociation constants (*Kd*s) i.e., tight binding affinities for all three putative AW boxes with *Kd* values between 0.27 and 5.24 nM, comparable with the affinity of WRI1 for the bona fide AW box1 from the promoter of the WRI1 target gene *BCCP2* of 0.65 nM (Maeo et al., 2009). Among the three BADC genes, WRI1 showed the highest affinity for the BADC1 AW box at 0.27 ± 0.27 nM. The AW box from the *BCCP1* promoter was also tested because although *BCCP1* contains an AW box (−60/-47 relative to TIS) the gene was shown not to be regulated by WRI1 (Fukuda et al., 2013). Consistent with this, the *Kd* value for dsDNA that includes the *BCCP1* AW box and WRI1 showed an approximately 37-fold higher *Kd* value than those of for *BCCP2* and WRI1 (SFig. 4). To test whether the consensus sequence [CnTnG](n)7[CG] in BADCs AW boxes is critical for WRI1 binding, 3 different mutant sequences were generated for each *BADC* AW box (Table S2) and tested in MST. As shown in Table S3, mutations in conserved nucleotides dramatically decreased their affinities to WRI1. Mutations in the 7 nucleotides between also decreased the affinity but to a less degree (Table S3).

**Figure 3.**
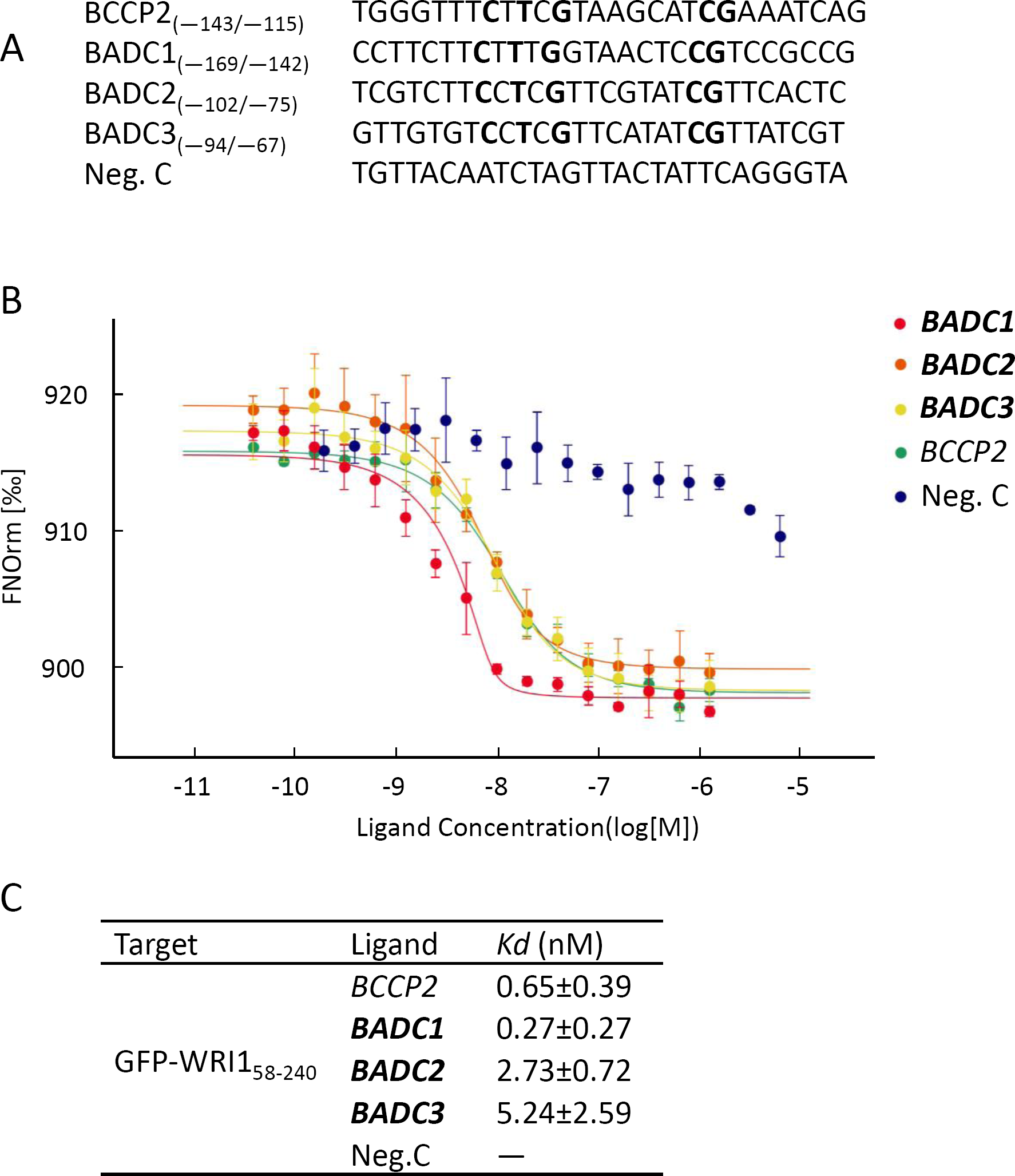
Purified WRI1 shows specific binding with AW boxes in the promoters of BADCs. In the thermophoresis experiment, A 28bp-oligonucleotide containing putative AW box for each gene (A) was titrated against purified WRI1 DNA binding domain (AA58-240). Dissociation curves for each pair are shown (B) and fitted to the data for calculation of the *Kd* values (C). An AW box in the promoter of *BCCP2* was used as a positive control. A random sequence is used as a negative control (Neg.C). AW box consensus sequences are marked in bold. Data shown are the mean±SD, n=3 independent repetitions.

**Figure 4.**
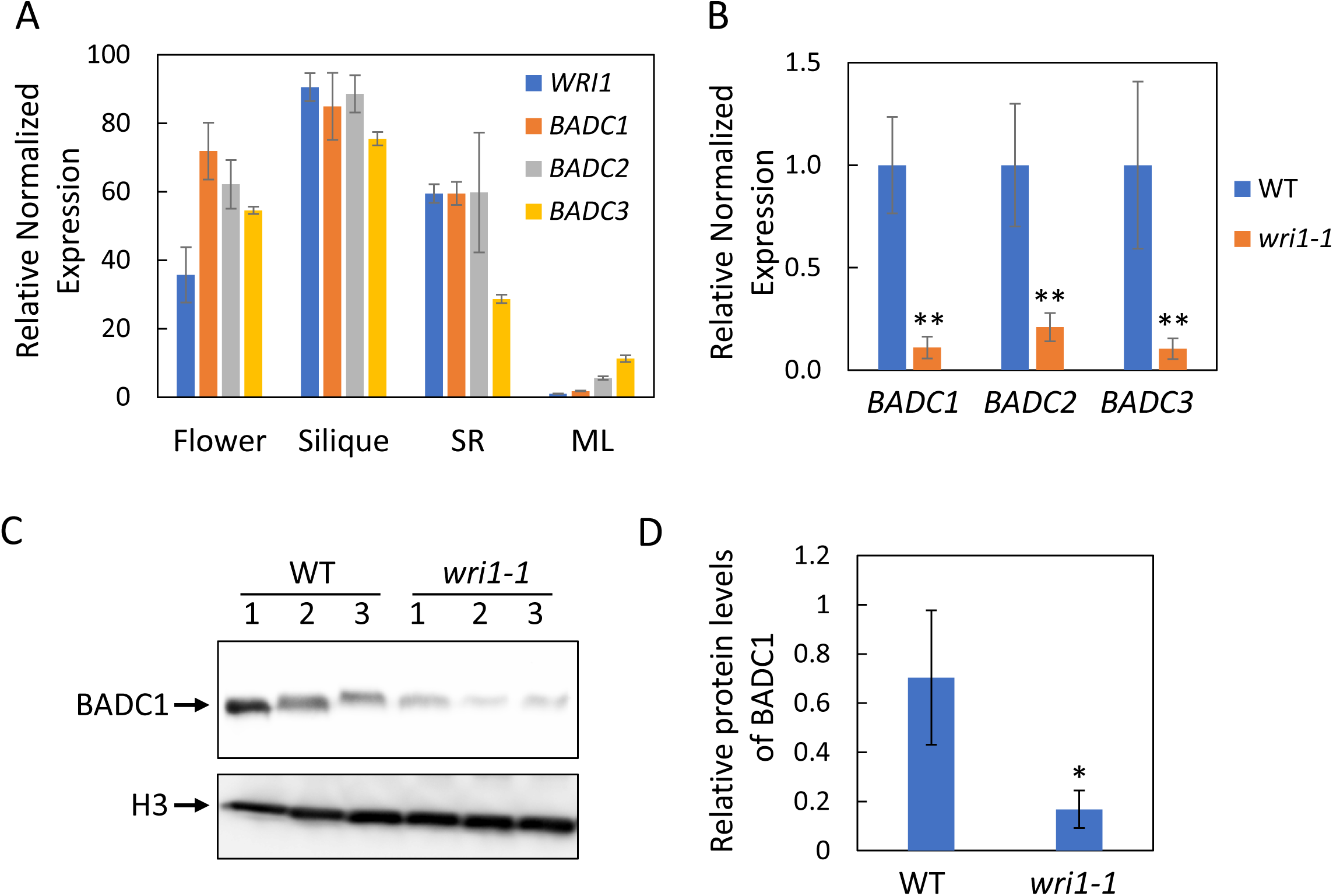
*BADCs* expression level and BADC1 protein level are significantly lower in *wri1-1* mutant developing siliques than in WT. (A) Tissue expression pattern of *WRI1* and *BADCs* in flowers, siliques (7 days after flowering), seedling roots (SR, 10 days old) and mature leaves (ML). (B) RT-qPCR results of *BADC1, BADC2*, and *BADC3* expression levels in siliques (7 days after flowering) of WT and *wri1-1* mutant. Values are means±SD from three independent experiments. Expression of each gene in WT sets to 1. For each experiment, total RNA was isolated from pooled siliques from WT and *wri1-1* plants. *Fbox* and *UBQ10* were used as reference genes. Asterisk denotes statistically significant difference from WT (using mean crossing point deviation analysis computed by the relative expression (REST) software algorithm, **, P<0.01). (C) BADC1 protein levels in siliques (7 days after flowering) of WT or *wri1-1* mutant are shown by immunoblot with BADC1 specific antibody. Siliques were collected from 3 independent plants of WT or *wri1-1* mutant. Protein loading is shown by histone H3 in the same protein samples. (D) The histogram shows relative BADC1 protein levels in (C) quantified with GelAnalyzer2010 and normalized against corresponding protein levels of histone H3. Asterisk denotes statistically significant difference from the WT (Student’s T test, *, P<0.05). Data in this figure is a representative of 3 independent repetitions.

### *BADCs* gene expression and BADC1 protein levels are lower in developing siliques of *wri1-1* than those of WT

Gene expression quantification of *WRI1* and *BADCs* in different tissues (flower, silique, root and mature leaf) showed generally similar tissue-specific expression pattern to that of *WRI1* (Fig. 4A), i.e., highly expressed in siliques (developing seeds) and low expression in mature leaves. Because of the highest expression of *WRI1* and *BADCs* in developing seeds among analyzed tissues (Fig. 4A), siliques were selected for investigating the relationship between *WRI1* and *BADCs*. Gene expression of *BADCs* in developing siliques (7 days after flowering) of *wri1-1* were only 10-20% of that in WT (Fig. 4B). We next used a BADC1 antibody that showed high specificity towards BADC1 protein as (Sfig.5) to probe BADC1 levels in *wri1-1*. Consistent with the observed decreased expression of *BADC1* in *wri1-1*, relative to WT, the level of BADC1 protein in *wri1-1* observed by immunoblotting was only 23% of that in WT (Fig. 4C). Since relative high gene expressions of *WRI1* and *BADCs* were also observed in seedling roots (Fig. 4A), BADC1 protein level was quantified in seedling roots and found to be significantly lower in *wri1-1* seedlings roots (12 day old) compared with wild type (Fig. S6).

### *BADCs* gene expression and BADC1 protein levels are significantly higher in developing siliques of *WRI1* overexpression transgenic plant than those of WT

To test whether overexpression of *WRI1* would increase gene expression of *BADCs* and the abundance of BADC1 polypeptide, two ethanol-inducible *WRI1* transgenic lines that were generated in our previous study (Zhai et al., 2017) were used. Both lines demonstrated elevated WRI1 accumulation after induction with 2% ethanol treatment (Zhai et al., 2017). As shown in figure 5A, *BADCs, WRI1* and *BCCP2* gene expression in developing siliques of *WRI1* transgenic plants that were induced by irrigation with 2% ethanol for 4 days were significantly higher than either WT or the corresponding *WRI1* transgenic plant lines that were not exposed to ethanol treatment (Fig. 5A). Consistent with the elevated gene expressions of *BADC1*, BADC1 protein levels were also significantly higher upon WRI1 overexpression (Fig. 5B and 5C). To test whether global regulation of WRI1 on BADCs, gene expression and protein level were also quantified in the seedling roots of *WRI1* transgenics, as shown in Fig. S7, both gene expression and protein level were higher in *WRI1* transgenics induced by ethanol treatment than either non-ethanol treatment control or WT (Fig. S7).

**Figure 5.**
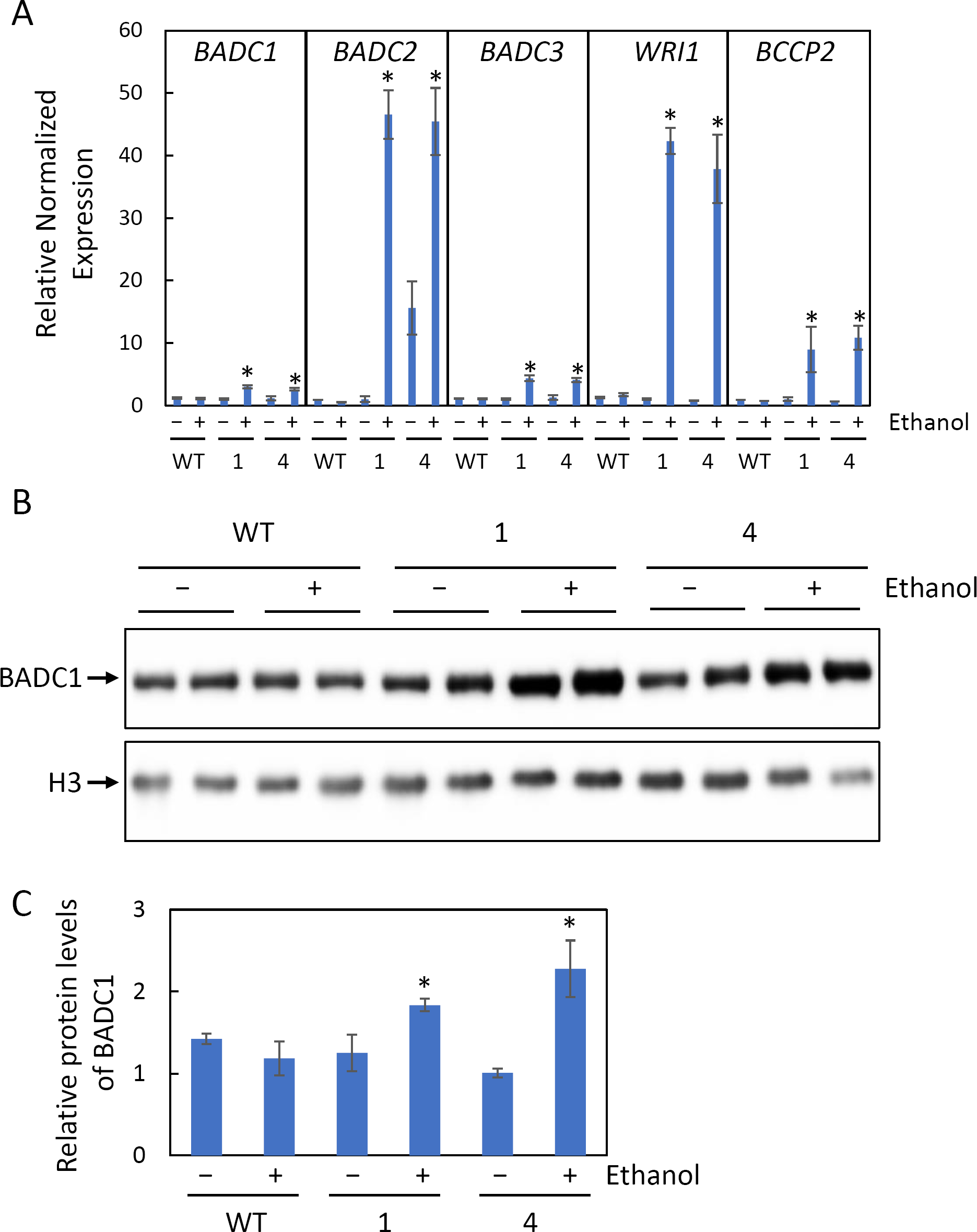
*BADCs* gene expression and BADC1 protein levels are elevated in inducible *WRI1* transgenic plants. (A) Relative gene expression of *BADCs, BCCP2* and *WRI1* in siliques (7 days after flowering) of WT or 2 independent ethanol inducible *WRI1* transgenic lines (AlcA:*WRI1* line1 and line4) treated (+) or not treated (−) with 2% ethanol for 4 days. (B) BADC1 protein levels. (C) The histogram shows relative BADC1 protein levels in (B) quantified with GelAnalyzer2010 and normalized against corresponding protein levels of histone H3. Asterisk denotes statistically significant difference from the WT (Student’s T test, *, P<0.05). Data in this figure is a representative of 3 independent repetitions.

### Overexpression of *BADC1* in *wri1-1* partially rescues *wri1-1* short-root phenotype

The data presented above supports the hypothesis that the expression of *BADC* genes are under the control of WRI1, and that the short-root phenotype of *wri1-1* primarily results from a deficiency of *BADC1* and *BADC2*. To test this hypothesis, the CDS of *BADC1* was placed under the control of the 35S promoter and transformed into *wri1-1*. 15 out of 17 independent transgenics demonstrated significantly longer primary root length than that of *wri1-1* when germinated and grown vertically on ½ strength MS medium plates for 16 days (Fig. 6A and 6B). Immunoblot showed that BADC1 protein levels were significantly increased in those *BADC1/wri1-1* transgenics (Fig. S8). To test whether IAA-Asp is somehow associated with the short-root phenotype of wri1-1, IAA contents were measured in 7-day-old seedlings of *BADC1/wri1-1*, *wri1-1* and WT and showed decreasing in *BADC1/wri1-1* when compared with *wri1-1*(Fig. 6C), supporting BADC1 being a downstream component of WRI1-dependent root development.

**Figure 6.**
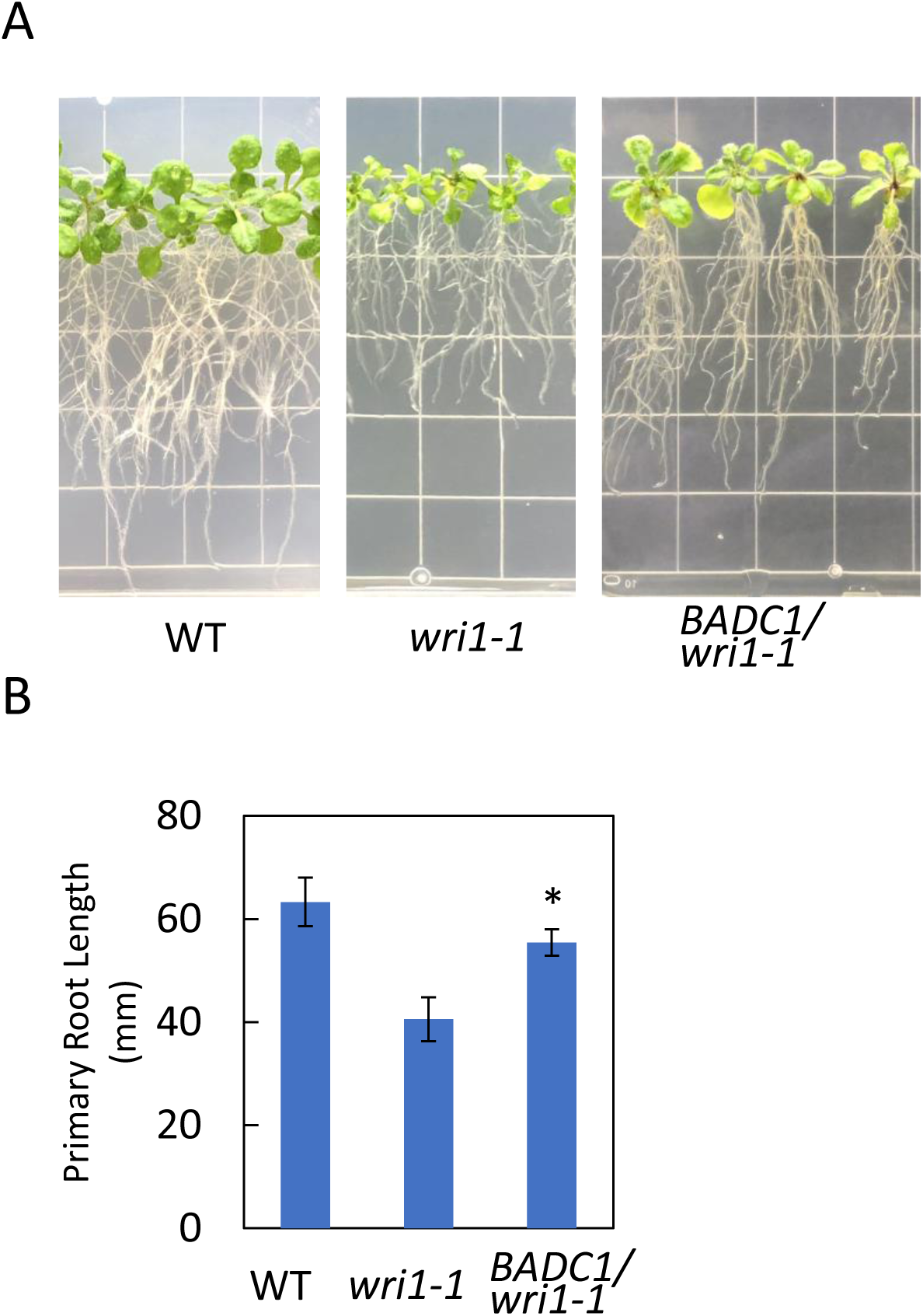
Overexpression of *BADC1* in *wri1-1* partially rescues *wri1-1* short-root phenotype. (A) Roots growth and B) Primary roots length measurement of 16-day-old WT, *wri1-1* and representative transgenic *wri1-1* overexpressing *BADC1* lines. Values in (B) are means±SD from 10 individual plants for each indicated genotype. Asterisk denotes statistically significant difference from *wri1-1* mutant (Student’s T test, *, P<0.05). (C) IAA-Asp was quantified in the 7-day-old seedlings of WT, *wri1-1* and *BADC1/wri1-1* transgenic line 1 and 2. Asterisk denotes statistically significant difference from *wri1-1* mutant (Student’s T test, *, P < 0.05). Data in this figure is a representative of 3 independent repetitions.

## Discussion

In this work we establish that expression of *BADC1, BADC2* and *BADC3* genes are under transcriptional control of WRI1. Evidence to support this comes from: 1) the identification of AW boxes within 200bp upstream of the TIS; 2) that microscale thermophoresis measurement demonstrated tight binding,i.e., *Kd*s in the low nanomolar range between WRI1 and the AW boxes from all three *BADC* genes; 3) that expression of each of the three *BADC* transcripts are reduced (by approximately 85%) in the *wri1-1* mutant; 4) the protein levels of BADC1 are reduced similarly to reductions in the level of its transcript in the *wri1-1* mutant. 5) Both gene expressions of *BADCs* and protein Levels of BADC1 are increased upon overexpression of *WRI1*.

That WRI1, the master transcriptional activator of genes involved in FA synthesis, simultaneously activate the transcription of BADCs which have been shown to act as negative regulators of ACCase might at first appear paradoxical. However, FA synthesis is metabolically expensive in terms of energy and reducing power, creating an imperative for tight metabolic control. For example, ACCase, the rate limiting step in FA synthesis is under tight genetic and biochemical control (Salie et al., 2016). When free fatty acid levels in the cell were increased beyond demand by supplementation with Tween-esters, two phases of feedback inhibition were reported, one short term, the second longer term (Andre et al., 2012). Short term inhibition occurs when the levels of oleoyl-ACP, the terminal product of the ACP-track of FA synthesis in the plastid, increase. Oleoyl-ACP inhibits ACCase by binding to it in a reversible manner (Andre et al., 2012). If, however, excess fatty acid levels persist for two or more days, an irreversible phase of inhibition of ACCase is initiated that is primarily mediated by BADC1 and BADD3 (Keereetaweep et al., 2018). In this context, BADC subunits should not be regarded as a simple inhibitor, but rather as a conditional inhibitor of ACCase. Thus, coexpression of the approximately 20 genes that promote FA synthesis along with the three BADC subunits makes biological sense because as WRI1 increases cellular metabolic demand by activating FA synthesis, it concomitantly increases the capacity to downregulate ACCase, and thereby FA synthesis, by increasing abundance of the conditionally inhibitory BADC subunits. Co-regulation of FAS and the BADCs thus allows the cell to match synthetic capacity with its regulatory capacity. The biochemical regulation of ACCase described herein is complementary to the transcriptional regulation of FAS we recently reported in which levels of the WRI1 polypeptide are tightly coupled to the availability cellular carbon in the form of sugars. When the carbon/energy status of the cell decreases, the SnRK1 carbon-sensing kinase becomes activated and it phosphorylates WRI1, leading to its selective proteasomal degradation thereby reducing activation of FAS genes (Zhai et al., 2017). Conversely, when carbon/energy abundance increases, levels of the SnRK1 kinase inhibitor trehalose 6-phosphatae increase, reducing WRI1 phosphorylation/degradation, thereby increasing the amount of WRI1 available to activate the transcription of FAS genes (Zhai et al., 2018).

In this study we also observed the *badc1 badc2* double mutant exhibits reduced primary root growth, a phenotype that resembled that of the *wri1-1* mutant (Kong et al., 2017), which in combination of our finding that WRI1 controls BADC transcription led us to ask whether the short-root phenotype characteristic of both *wri1-1* and *badc1badc2* mutants result from a deficit of BADC. If correct, overexpression of BADC in *wri1-1* would mitigate the *wri1-1* short-root phenotype. This view is supported by the observations that overexpression of *BADC1* in *wri1-1* partially rescued its short-root phenotype in a manner similar to its overexpression in the *badc1badc2* mutant background (and as overexpression of *BADC2* had in the *badc1badc2* mutant background) and reduced IAA-Asp content in *BADC1/wri1-1*. Taken together this work suggests that deficiency of BADC levels, resulting either directly via mutation of *badc* genes, or indirectly through the mutation of their upstream transcriptional activator, *wri1*, result in a short-root phenotype *via* an as yet-to-be determined mechanism. Further work will be required to identify details of the mechanism, that is likely related to increased levels of conjugated IAA that were observed for both *wri1-1* and *badc1badc2* mutants.

Taken together, we show all three BADC genes are direct target genes of WRI1 and can play a role in root development. The observation that WRI1 not only upregulates genes contributing to FA synthesis, but also BADCs which are conditional regulators of FA synthesis identifies an additional layer of regulation with respect to FAs synthesis.

## Materials and Methods

### Plant Materials and Growth Conditions

Wild-type (WT) *Arabidopsis thaliana* (ecotype Columbia-0) and *wri1-1*(CS69538), *badc1* (Salk 000817C), *badc2* (Salk 021108C) and *badc3* (CS2103834) were used in this study. Double mutant *badc1 badc2* was generated and described previously (Keereetaweep et al., 2018). For growth on plates, seeds were surface-sterilized with 70% ethanol, followed by 30% bleach containing 0.01% Tween20, and rinsed three times with sterile water. Seeds were stratified for 3 days at 4°C in the dark and germinated on half Murashige and Skoog (MS) medium supplemented with 1% w/v sucrose plates in an incubator with a light/dark cycle of 18h/6h at 23°C, photosynthetic photon flux density of 250 μmol m^−2^ s^−1^, and 75% relative humidity.

### Genetic Constructs

CDS of *BADC1* and *BADC2* were amplified by PCR from cDNAs using primers listed in Table S4. The PCR products were then cloned into the Invitrogen GATEWAY^M^ pDONR/Zeo vector (Thermo Fisher Scientific, Waltham, MA; www.thermofisher.com) using the BP reaction and sub-cloned (LR reaction) into the plant GATEWAY™ binary vector: pGWB414 (Nakagawa et al., 2007). *BADC1*/pGWB414 and *BADC2*/pGWB414 were introduced into *badc1 badc2* double mutant by agrobacterium (AgL0)-mediated transformation respectively. For rescuing short-root phenotype of *wri1-1, BADC1*/pGWB414 was transformed into *wri1-1* mutant.

### Quantification of IAA, IAA-Asp in badc1 badc2 double mutant Seedlings

7-day-old Arabidopsis WT and *badc1 badc2* seedlings grown vertically on half MS medium supplemented with 1% w/v sucrose were harvested (20-30 seedlings per replicate, ∼100 mg FW) and frozen in liquid Nitrogen. Frozen samples were sent to the Proteomics & Mass Spectrometry Facility at the Donald Danforth Plant Science Center for acidic hormone analysis where frozen plant material is extracted twice with ice-cold acetonitrile/methanol (1:1 v:v) using a bead beater and resulting samples were dried with the use of a speedvac. Samples were reconstituted in 30% MeOH and separated via high performance liquid chromatography using a Waters (Milford, MA) Acquity UPLC® BEH C18 1.0 × 100 mm, 1.7 μm column kept at 50 °C with a flow rate of 15 μL/min coupled to a a Sciex 6500 QTrap® mass spectrometer (Framingham, MA) operated in multiple reaction monitoring (MRM) mode employing polarity-switching for simultaneous detection of positive and negative ions. Data analysis was completed using MultiQuant 3.0.2 (AB Sciex) with unlabeled compound peak areas normalized to their respective labeled internal standard peak areas. Labeled standards for indole-3-acetic acid and N-(3-indolylacetyl)-DL-aspartic acid were obtained from Sigma-Aldrich. Calibration curves were linear (*r* values = >0.99) within the ranges provided above and applying 1/x weighting.

### RNA Isolation and RT-qPCR

To quantify gene expression in mutants and transgenic plants, total RNAs were isolated using the RNeasy Plant Mini Kit (Qiaqen, Gaithersburg, MD). cDNA was prepared using SuperScript™ IV VILO™ Master Mix with ezDNase™ Enzyme (Invitrogen, Carlsbad, CA). RT-qPCR reactions were setup with SsoAdvanced™ Universal SYBR^®^ Green Supermix (Bio-Rad) and gene specific primers (primer sequences are listed in Table S4) and performed with the CFX96 qPCR Detection System (Bio-Rad). Statistical analysis of qRT-PCR data was carried out with REST2009.

### Expression of purification of recombinant GFP-WRI1_58-240_

DNA sequence of full length of WRI1 fused with GFP on its N-terminus (*GFP-WRI1*) was amplified from a OWD5 vector previously described in Zhai et al. (Zhai et al., 2017). *GFP-WRI1* then was inserted into pet28b between *XhoI* and *NcoI* by in-fusion cloning. Full length of *WRI1* in the resulted *GFP-WRI1*/pet28b was replaced with DNA sequence corresponding to WRI1 DNA binding domain (WRI1_58-240_) between *XhoI* and *BsrGI* (Kong et al., 2017). The primer pairs used in building this construct is listed in Table S4. Recombinant GFP-WRI1_58-240_ was expressed in *E.coli* BL21(DE3). Protein purification was performed as reported by Zhai (Zhai et al., 2017).

### MicroScale Thermophoresis (MST)

Thermophoretic assays were conducted using a Monolith NT.115 apparatus (NanoTemperTechnologies, South San Francisco, CA; nanotempertech.com). For determining *Kd*s, 8 nM of GFP-WRI1_58-240_ was incubated with a serial dilution of the ligand (dsDNA) from 1.25 μM to 38.81 pM. Samples of approximately 10 μl were loaded into capillaries and inserted into the MST instrument loading tray (Monolith NT.115). The thermophoresis experiments were carried out using 40% MST power and 80% LED power at 25 °C.

### Antibody and Immunoblotting

Anti-BADC1 and anti-histone H3 antibodies were used in this study. Anti-BADC1 antibodies were requested from Dr. Jay Thelen (Salie et al., 2016). Histone H3 polyclonal antibodies were purchased from Agrisera (Catalog No. AS10710, Vännäs, Sweden; www.agrisera.com). A total of 50 mg of siliques (7 days after flowering, DAF) of WT and *wri1-1* mutant plants were ground in liquid nitrogen and then mixed with 200 mL of preheated protein extraction buffer (8 M urea, 2% SDS, 0.1 M DTT, 20% glycerol, 0.1 M Tris-HCl, pH (6.8), and 0.004% Bromophenol Blue). Samples were centrifuged at 17,000 g before transferring the supernatant to a new microcentrifuge tube and then loading into SDS-PAGE. 20 ul of supernatant was loaded each and resolved in SDS-PAGE and transferred onto PVDC membrane for immunoblot analysis. Primary antibodies anti-BADC1 were used at a 1:3000 dilution, and the anti-H3 antibodies were used at a 1: 5000 dilutions. Immunoblots of targeted proteins were visualized using HRP-conjugated secondary antibodies (Catalog No. AP187P, Millipore) with SuperSignal™ West Femto Maximum Sensitivity Substrate (Catalog No. 34095, ThermoFisher). Immunoblot signals were detected and digitalized with Image Quant LAS4000 and quantified with GelAnalyzer2010a.

## List of author contributions

H.L, Z.Z. J.Sc. J.K., and J.Sh. conceived the original research plans and designed the experiments; H.L, K.K., Z.Z. and J.K. performed the experiments; H.L, Z.Z. J.Sc. J.K., and J.Sh. analyzed the data and wrote the article with contributions of all the authors; J.Sh. agrees to serve as the author responsible for contact and ensures communication.

## ACCESSION NUMBERS

Sequence data from this article can be found in The Arabidopsis Information Resource under the following accession numbers: *WRI1* (AT3G54320), *BADC1* (AT3G56130), *BADC2* (AT1G52670), *BADC3* (AT3G15690), *BCCP1* (AT5G16390), *BCCP2* (AT5G15530), *Fbox* (AT5G15710) and *UBQ10* (AT4G05320).

## Supplemental Figure Legends

**Supplemental Figure 1**. *BADC1* expression in transgenic *badc1badc2* double mutant overexpressing *BADC1* lines.

**Supplemental Figure 2**. *badc1badc2* shows similar short-root phenotype as *wri1-1.*

**Supplemental Figure 3** Purified recombinant GFP-WRI1_58-240_.

**Supplemental Figure 4**. Putative AW boxes show varied binding affinity with WRI1.

**Supplemental Figure 5**. The BADC1 antibody used in this study specifically recognizes BADC1.

**Supplemental Figure 6**. BADC1 protein levels are lower in the roots of seedlings of *wri1-1* than WT. **Supplemental Figure 7**. Both *BADC1* expression and protein levels are elevated in the roots of *WRI1* inducible expression transgenic plants.

**Supplemental Figure 8**. BADC1 protein levels in the *BADC1*/*wri1-1* transgenic line.

**Table S1.** Quantification of plant hormones in *badc1 badc2* double mutant seedlings.

|  | ABA | JA | SA | OPDA | JA-Ile | t-Z riboside |
| --- | --- | --- | --- | --- | --- | --- |
| WT | 2.80±0.45 | 12.8±4.15 | 21.06±1.30 | 238.82±109.75 | 0.5±0.31 | 2.63±0.93 |
| *badc1 badc2* | 3.33±0.53 | 12.2±7.35 | 26.36±7.43 | 285.13±140.5 | 0.35±0.31 | 1.96±1.90 |

Plant hormones concentration are shown as ng per g DW in this table.

**Table S2.**
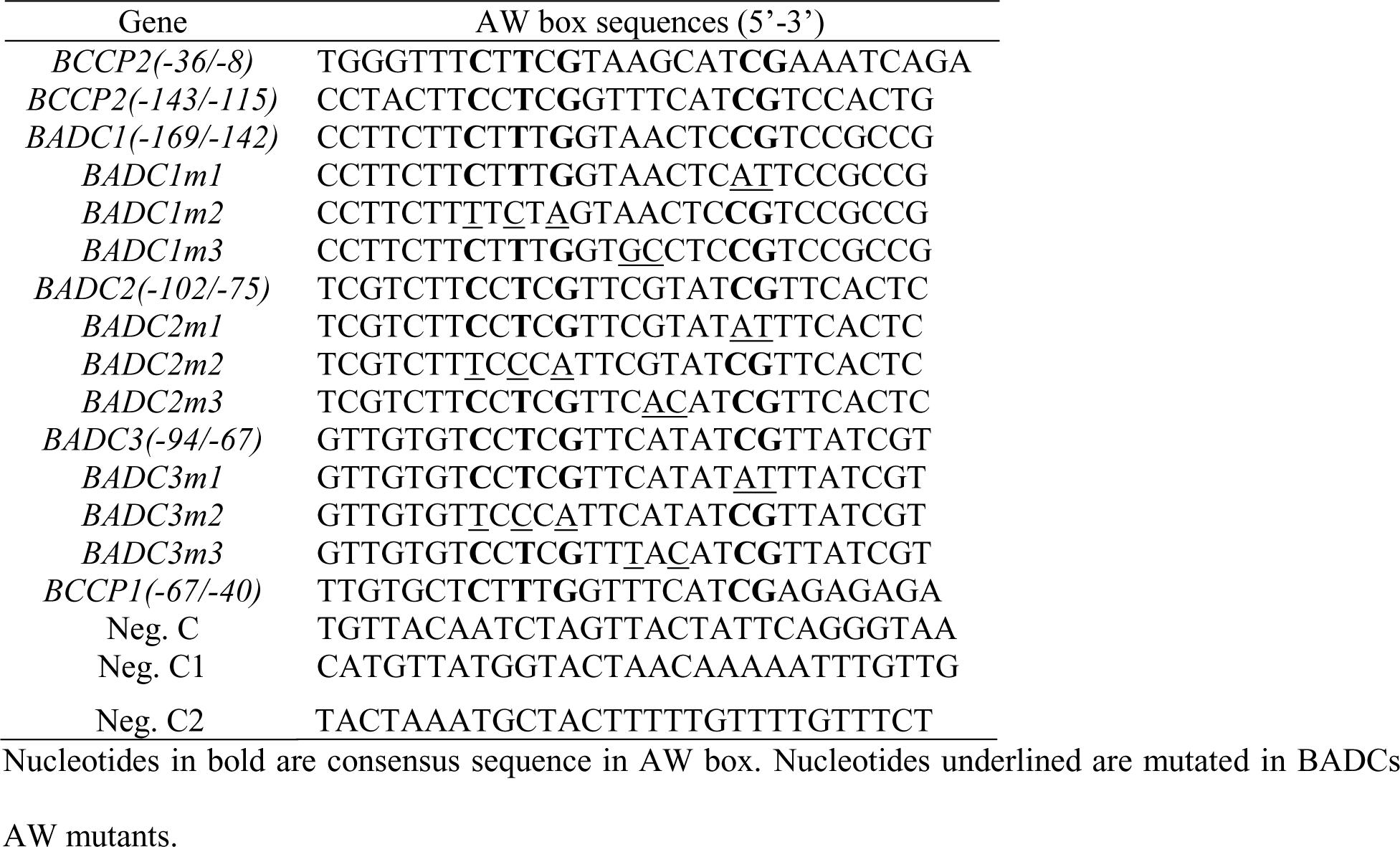
Nucleotide sequences of AW box in thermophoretic experiment.

**Table S3.** Binding affinity of *BADCs* AW boxes and their mutated ones to WRI1 were quantified as equilibrium dissociation constants (*kd*) using microscale thermophoresis.

| AW boxex | *Kd* (nM) |
| --- | --- |
| *BADC1* | 0.27±0.27 |
| *BADC2* | 2.73±0.72 |
| *BADC3* | 5.24±2.59 |
| *BADC1m1* | > 1000 |
| *BADC2m1* | > 1000 |
| *BADC3m1* | > 1000 |
| *BADC1m2* | > 1000 |
| *BADC2m2* | > 1000 |
| *BADC3m2* | > 1000 |
| *BADC1m3* | 70.67 ± 27.80 |
| *BADC2m3* | 149.18 ± 138.63 |
| *BADC3m3* | 22.88 ±2.94 |

**Table S4.**
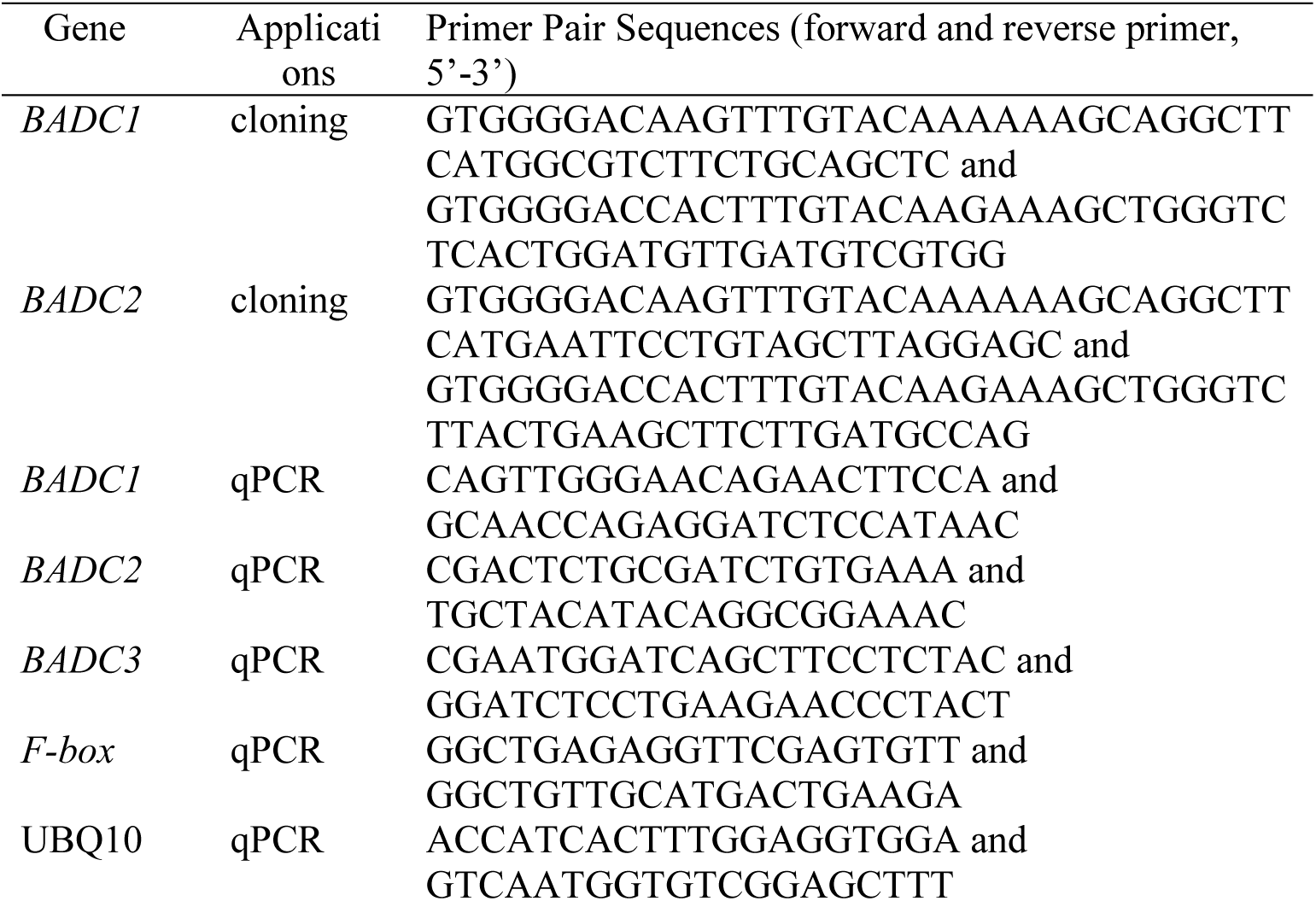
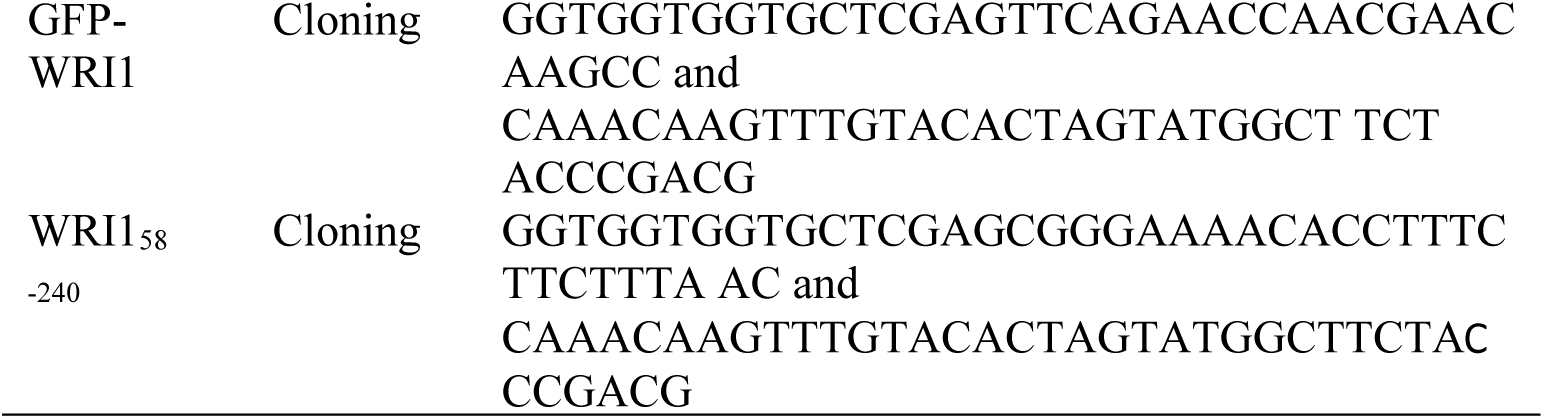
Oligonucleotide sequences of primers used in this study.

## Acknowledgements

We thank Dr. Jay Thelen (University of Missouri) for the gift of BADC1 antibody.

## Notes

This work was supported by the Division of Chemical Sciences, Geosciences, and Biosciences, Office of Basic Energy Sciences, US Department of Energy (grant DOE KC0304000 JSh) (and DE-SC0012704 JSc), and by a National Science Foundation under which initial discussions took place and several reagents were made (Grant DBI 1117680 JSh and JSc).

The authors declare no conflict of interest.

