## Supplemental Figure 1-8 for "BIOTIN ATTACHMENT DOMAIN-CONTAINING proteins, inhibitors of ACCase, are regulated by WRINKLED1"

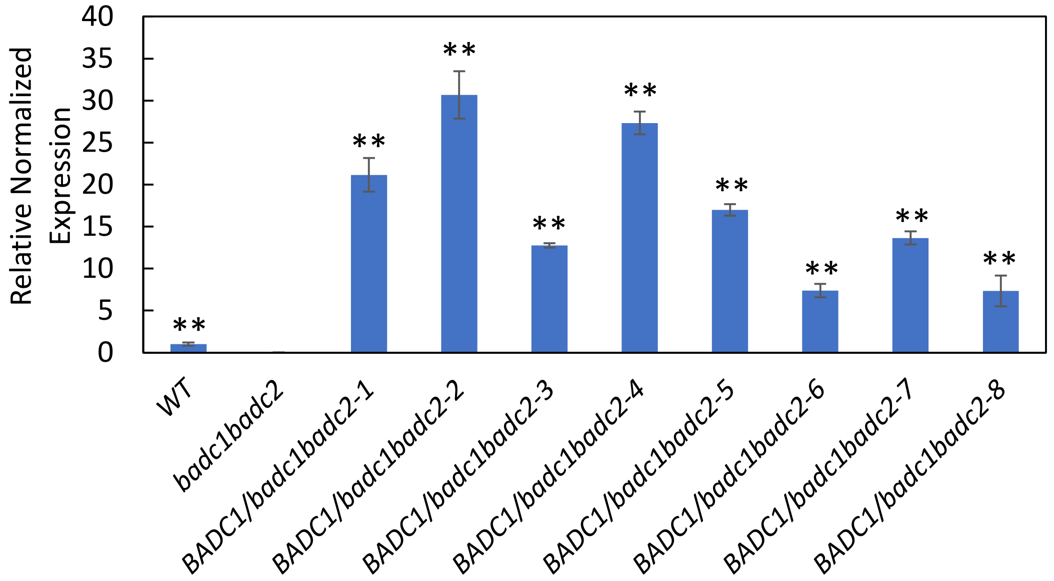


**Supplemental Figure 1**. *BADC1* expression in transgenic *badc1 badc2* double mutant overexpressing *BADC1* lines. RT-qPCR results of *BADC1* expression levels in 7-day-old seedlings of 8 independent transgenic lines. Values are means±SD from three independent experiments. Expression of *BADC1* in WT sets to 1. For each experiment, total RNA was isolated from pooled seedlings from indicated genotypes. Asterisk denotes statistically significant difference *from badc1 badc2* (using mean crossing point deviation analysis computed by the relative expression (REST) software algorithm, **, P<0.01).

A

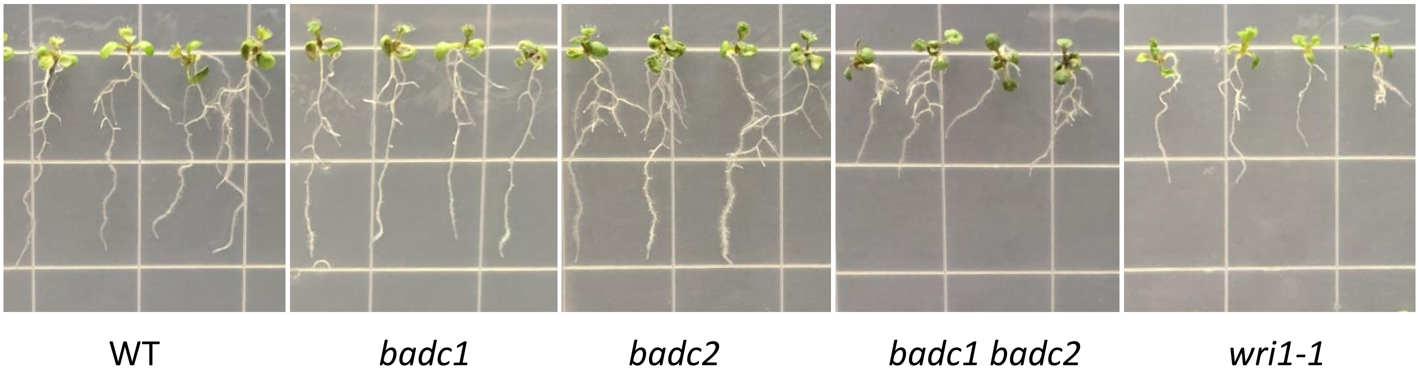


B

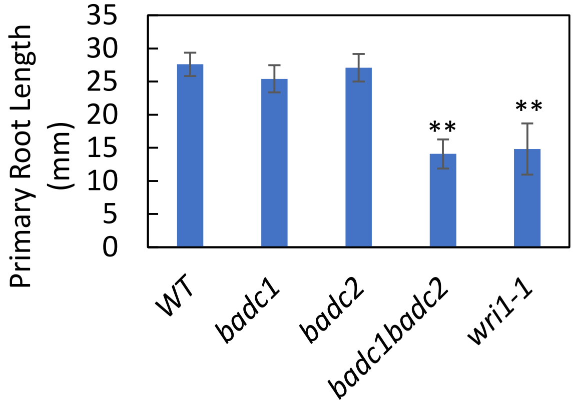


**Supplemental Figure 2**. *badc1 badc2* shows similar short-root phenotype as *wri1-1.* (A) WT, *badc1*, *badc2*, *badc1 badc2* and *wri1-1* germinated and grown vertically on 1/2MS media supplemented with 1% of sucrose for 7 days before photographed. *(*B) Primary root length was measured by ImageJ. Values are means±SD from 10 individual plants for each indicated genotype. Asterisk denotes statistically significant difference from the WT (Student's T test, **, $P<0.0001$).

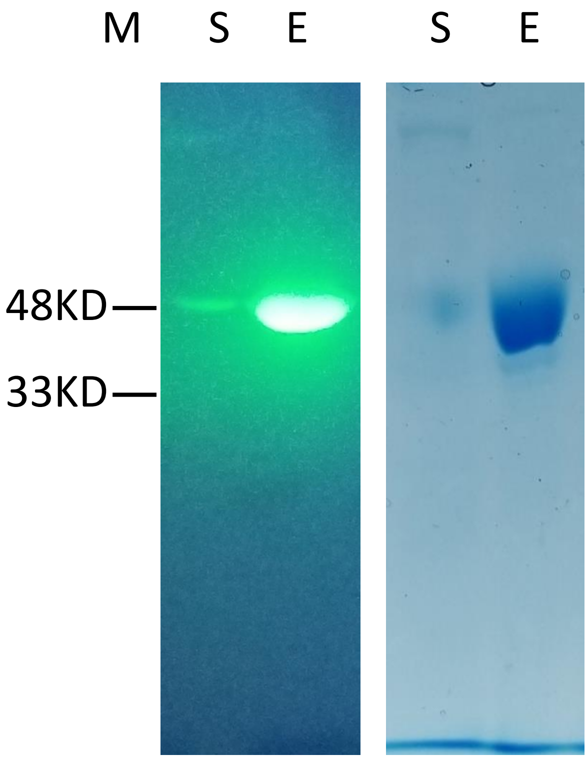


**Supplemental Figure 3**. Purified recombinant GFP-$WRI1_{58\text{-}240}$. HA-tagged GFP-$WRI1_{58\text{-}240}$ was expressed in *E.coli ,* bound with and eluted from Ni-NTA, mixed with non-denaturing loading buffer (no heat denaturing) and run on SDS-PAGE. GFP fluorescence under blue light (Left). Coomassie Blue staining (Right). M, protein marker. S, supernatant. E, elution.

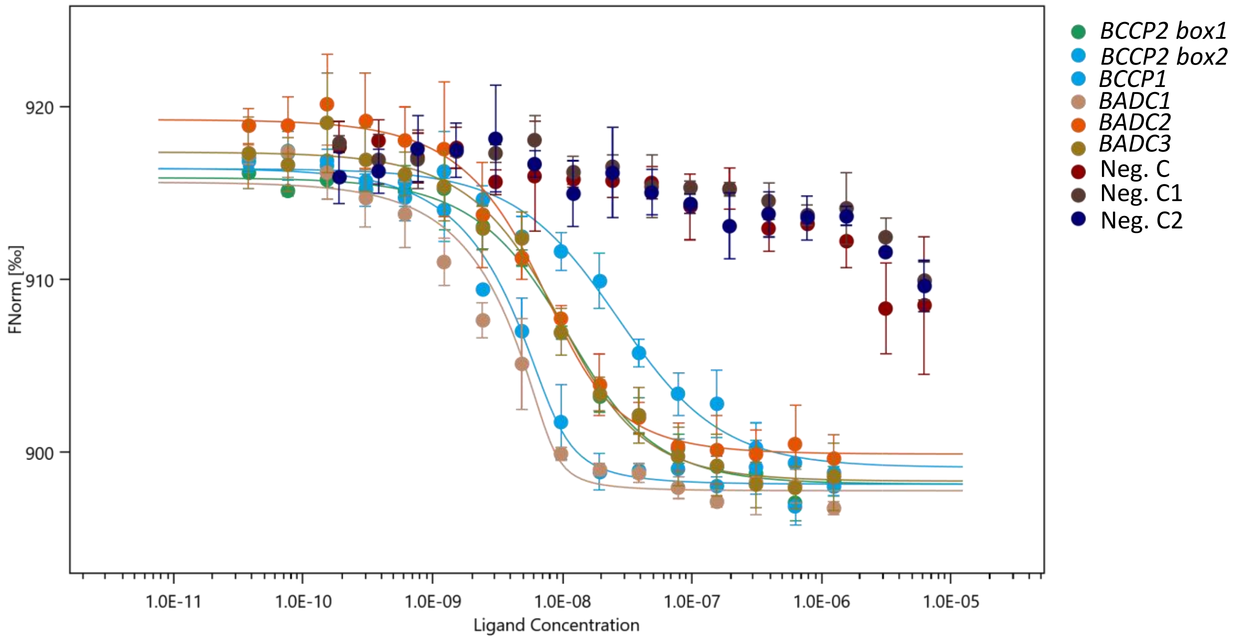


| Target | Ligand(s) | Kd (nM) |
|---|---|---|
| | BCCP2 box1 | 0.65±0.39 |
| | BCCP2 box2 | 6.68±0.94 |
| | BCCP1 | 23.85±4.42 |
| GFP-WRI1$_{58\text{-}240}$ | BADC1 | 0.27±0.27 |
| | BADC2 | 2.73±0.72 |
| | BADC3 | 5.24±2.59 |
| | Neg.C | — |
| | Neg.C1 | — |
| | Neg.C2 | — |

**Supplemental Figure 4**. Putative AW boxes show varied binding affinity with WRI1. In addition to pairs in Figure 3, BCCP1 AW box2, BCCP2 AW box and 2 more negative controls (Neg.C1 and Neg.C2) are also included in thermophoretic experiments to demonstrate varied binding affinity with WRI1. Neg.C1 and C2 are random DNA sequence. DNA sequences in this figure were shown in Table S2.

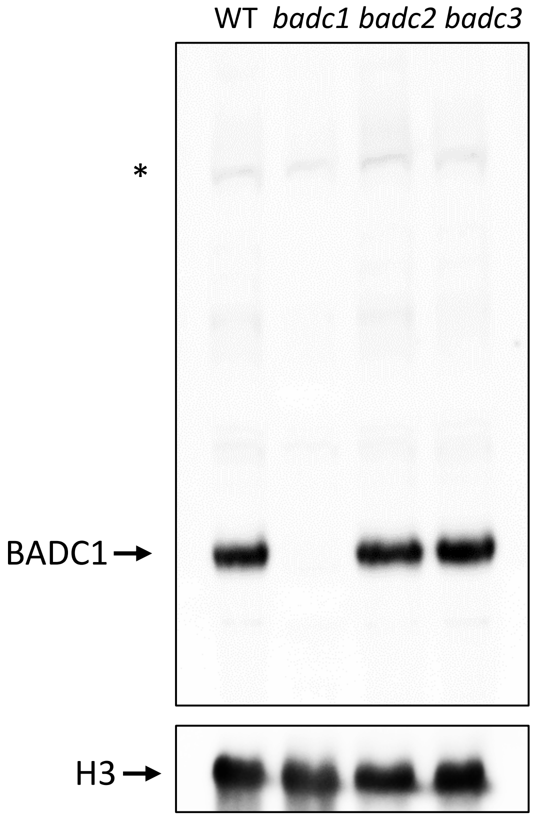


**Supplemental Figure 5**. The BADC1 antibody used in this study specifically recognizes BADC1. BADC1 protein levels in 7-day seedlings of WT or *badc* mutants (*badc1*, *badc2* and *badc3*) are shown by immunoblot with BADC1 antibody. Protein loading is shown by histone H3 (H3) in the same protein samples. Star indicates a non-specific band.

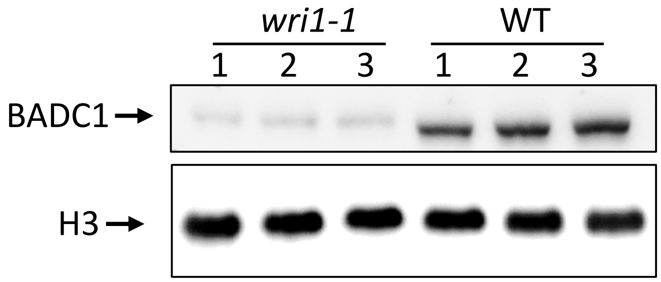


**Supplemental Figure 6**. BADC1 protein levels are lower in the roots of the seedlings of *wri1-1* than WT. BADC1 protein was quantified in the root tissues collected from 12-day-old seedlings of wild type and *wri1-1* mutant respectively. Protein loading is shown by histone H3 (H3) in the same protein samples.

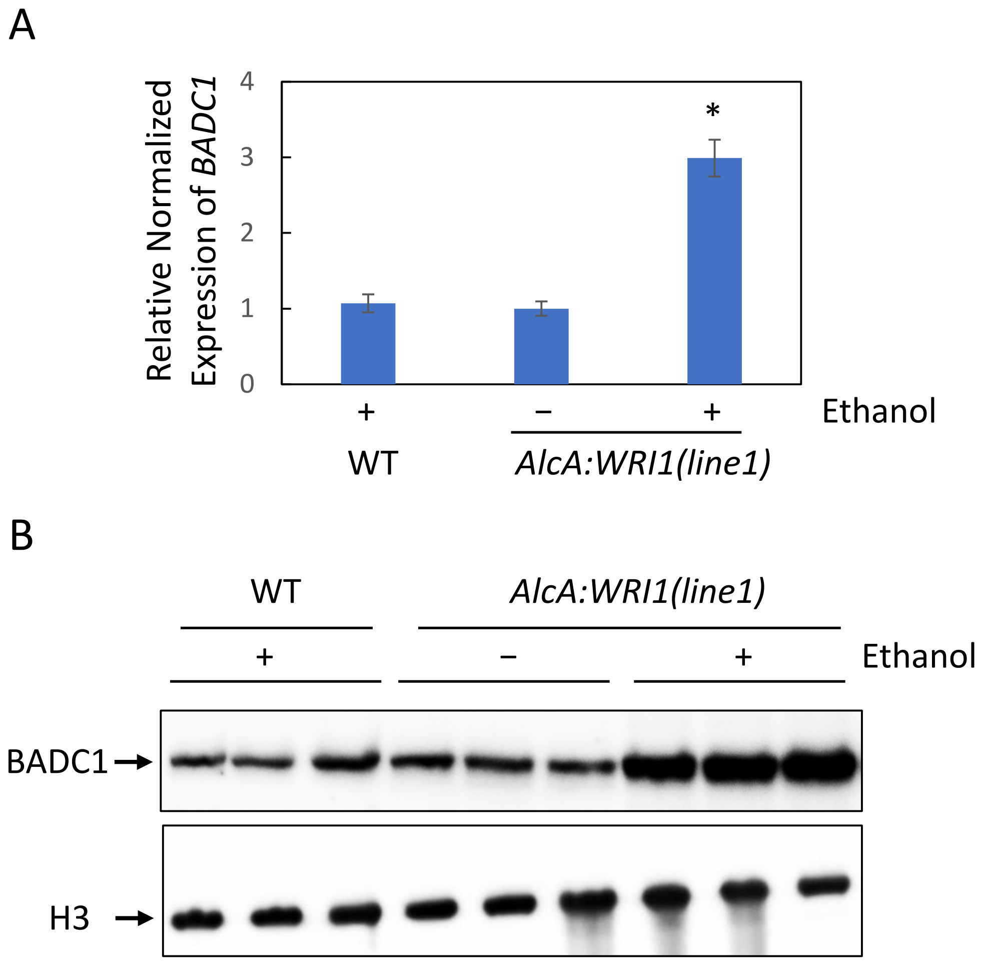


**Supplemental Figure 7**. Both *BADC1* expression and protein levels are elevated in the roots of *WRI1* inducible expression transgenic plants. (A) *BADC1* expression in the roots of 10-day-old seedlings of WT, ethanol inducible *WRI1* transgenic plant (line1) treated (+) or not treated (−) with 2% ethanol for 3 days. (B) Correspondingly, BADC1 protein levels were quantified by immunoblot with BADC1 specific antibody. Protein loading is shown by histone H3 in the same protein sample. Asterisk in this figure denotes statistically significant difference from the WT (Student's T test, *, $P<0.05$).

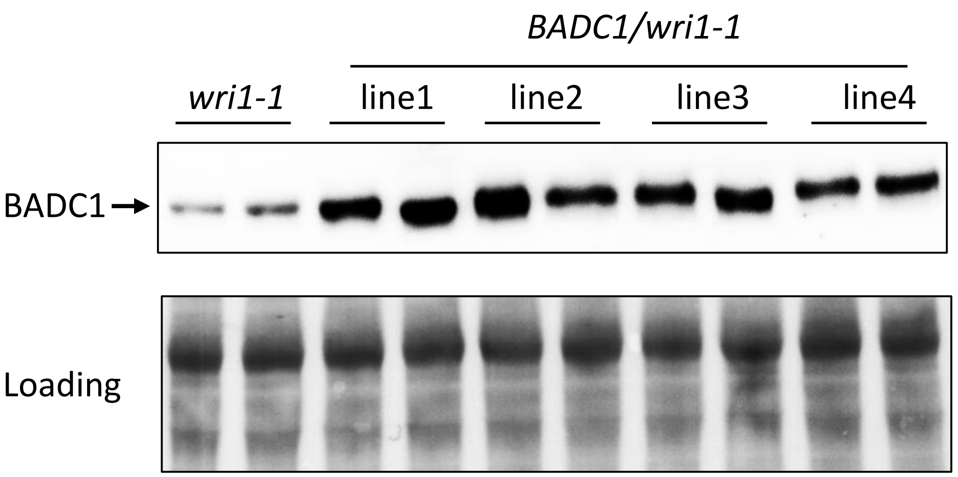


**Supplemental Figure 8**. BADC1 protein levels in the *BADC1*/*wri1-1* transgenic lines. BADC1 protein levels were quantified in the seedlings of 4 independent 10-day-old *BADC1/wri1-1* transgenic lines (line1-4) and of wild type. Protein loadings are shown by Ponceau staining with the same blot.
